# A Multi-Dimensional Approach to Map Disease Relationships Challenges Classical Disease Views

**DOI:** 10.1101/2024.02.15.580369

**Authors:** Lena Möbus, Angela Serra, Michele Fratello, Alisa Pavel, Antonio Federico, Dario Greco

## Abstract

The categorization of human diseases is mainly based on the affected organ system and phenotypic characteristics. This is limiting the view to the pathological manifestations, while it neglects mechanistic relationships that are crucial to develop therapeutic strategies. This work aims to advance the understanding of diseases and their relatedness beyond traditional phenotypic views. Hence, the similarity among 502 diseases is mapped using six different data dimensions encompassing molecular, clinical, and pharmacological information retrieved from public sources. Multiple distance measures and multi-view clustering is used to assess the patterns of disease relatedness. The integration of all six dimensions into a consensus map of disease relationships reveals a divergent disease view from the International Classification of Diseases (ICD), emphasizing novel insights offered by a multi-view disease map. Disease features such as genes, pathways, and chemicals that are enriched in distinct disease groups are identified. Finally, an evaluation of the top similar diseases of three candidate diseases common in the Western population shows concordance with known epidemiological associations and reveals rare features shared between Type 2 diabetes and Alzheimer disease. A revision of disease relationships holds promise for facilitating the reconstruction of comorbidity patterns, repurposing drugs, and advancing drug discovery in the future.

## 1 Introduction

Human diseases have traditionally been considered as phenotypes defined by their observable signs, symptoms, anatomical location of manifestation, and immediate trajectories. Despite the numerous molecular insights gained after the completion of the Human Genome Project, our current disease deffinitions are still mainly phenotype-based. Indeed, we group diseases by anatomical entities (*e.g.*, skin diseases, diseases of the circulatory system), and broad etiological categories (*e.g.*, cancerous, infectious, or congenital diseases) (*1*). This is practical in clinical routines to quickly narrow down the diagnostic options and streamline therapy schemes as well as for administrative purposes. However, this strategy neglects scenarios where (1) two diseases could have a common phenotype, but a fundamentally different mechanistic basis and where (2) two diseases co-occurring in a patient could share a mechanistic basis, although their phenotypic manifestations might be distinct (*2*). For the development of novel therapies, especially in the light of rare diseases, precision medicine and multimorbidity, mechanistic categorizations of diseases and eventually individual patients are needed (*3*, *4*).

A significant amount of disease-related data is currently publicly available, including high-throughput omics, experimental, and epidemiological data as well as inferred or summarized data from computational data integration approaches. Additionally, there are several systems organizing diseases such as the International Classification of Diseases (ICD) as well as medical ontologies organized in hierarchies or knowledge graphs (*5*, *6*). Multiple approaches to map diseases and examine their relationships based on different data dimensions have been performed. Disease relationships are modelled in networks, distance matrices or hierarchies and can be evaluated by network topological measures or distance metrics such as the distance of two diseases in a hierarchy or the distance based on overlapping characteristics (*7*). These methodologies suggested new or confirmed suspected disease associations such as between vitamin B deficiency and endogenous depression, breast and lung cancer, psoriasis and asthma as well as between diabetes and neoplasms and cerebral degenerative disorders (*8–10*). Few studies addressed the relationship of comorbidities by *e.g.*, inferring comorbidity links from patient histories (*11*). Modelling disease histories of patients by network analysis showed that diseases close in the network are more likely to become comorbid diseases (*12*). Together, these approaches aim to overcome the concept of phenotype-based disease definitions. Broadening our conventional understanding of diseases has the potential to elucidate and anticipate patterns of comorbidity, enhance drug reformulation, and catalyze drug discovery efforts.

Disease relationships have been explored based on one to three different data dimensions. Moreover, existing studies revealed that the omission of data layers decreased the performance of mapping diseases (*10*). A multi-dimensional view is needed to build a comprehensive mechanistic understanding of pathological phenotypes by combining multiple data layers generating a more informed and robust picture of disease similarities. Such types of consensus approaches seamlessly integrate complementary information, thus offering a comprehensive understanding of the intricate connections between diseases. At the same time, multi-view approaches help decreasing the impact of (unknown) biases by false or incomplete data in single dimensions or for single diseases. Here, we mapped the relationships of 502 diseases based on a consensus of six data dimensions including genomic, clinical, and chemical data. Beside an exploration of the frequency of disease-associated features, our main contributions are a map of disease-to-disease relationships based on single dimensions as well as their consensus, a comparison of this with the ICD-10 system, and an evaluation of the direct neighborhood of three common diseases in the Western population. We show that clustering diseases based on multiple dimensions breaks classical phenotypic disease views and reveals dominating features of new disease groups.

## 2 Results and Discussion

We developed a computational framework to study disease-to-disease similarities based on six data dimensions: (1) disease-associated genes, (2) disease-associated pathways, (3) chemicals medicating diseases, i*.e.*, drugs, (4) disease-associated symptoms, (5) chemicals targeting disease-associated genes, and (6) disease-associated chemicals. By combining clinical information such as symptoms and drugs with mechanistic information including genes, pathways, and chemicals, we gain a holistic view of disease landscapes. The data of the six dimensions were retrieved from a previously established knowledge space, as introduced by the authors. In this space, publicly available data have been curated, homogenized, and organized into a graph format to facilitate accessibility and analysis (*13*). In Figure 1, we visualize the six dimensions as they have been curated in our knowledge space including the original sources the data have been retrieved from.

**Figure 1:**
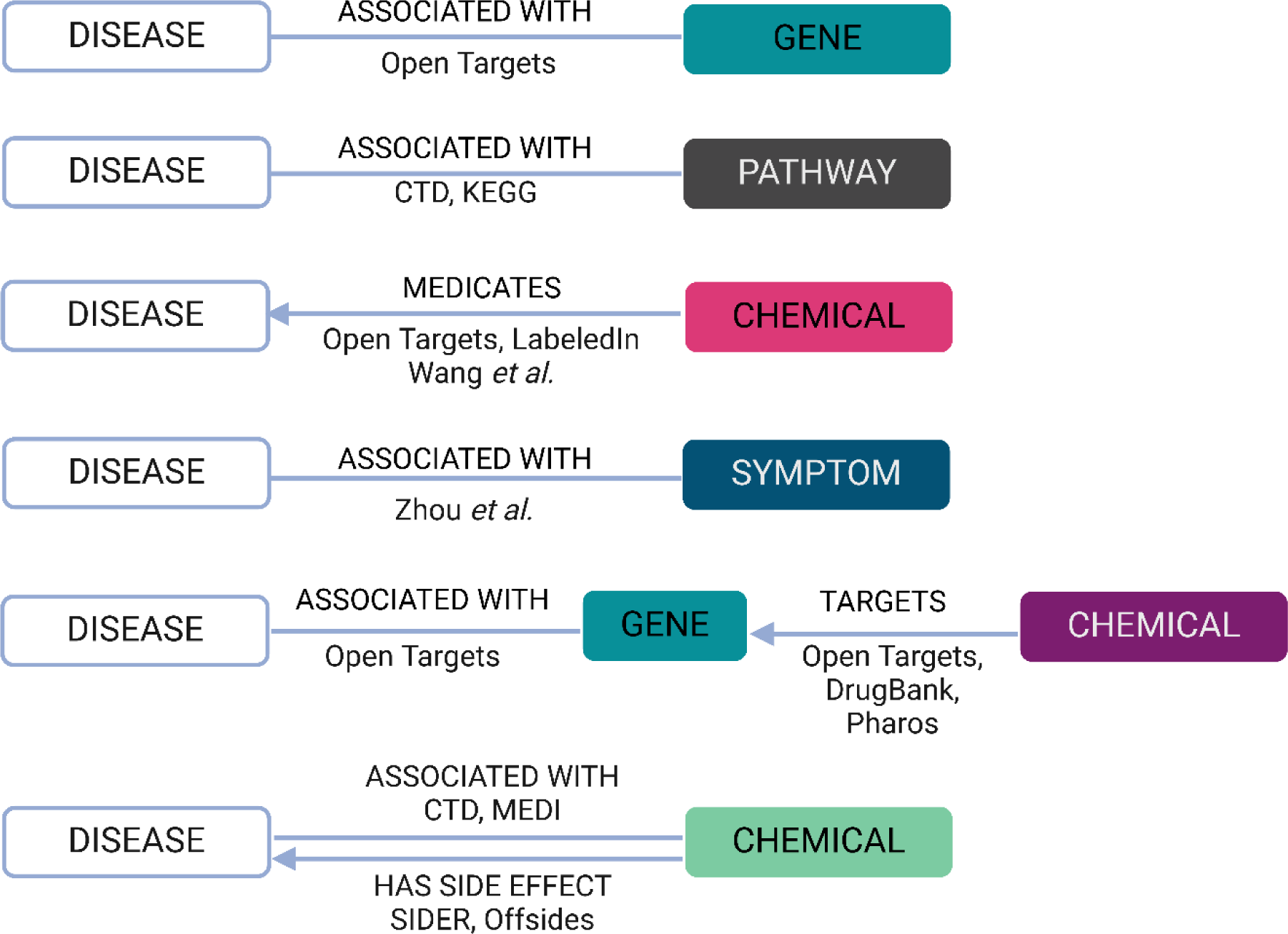
Six data dimensions for diseases. Disease-related data from six dimensions were retrieved from our internal knowledge base. Original data sources are indicated on the edges.

We retrieved data about 7,618 diseases with associated genes, 4,894 diseases with associated pathways, 1,811 diseases with medicating chemicals, 4,579 diseases with associated symptoms, 7,188 diseases with chemicals targeting associated genes, and 16,723 diseases with associated chemicals (Supporting Data 1). Hence, the number of diseases with available data varies considerably among the six dimensions. For the usability of data for the disease dimensions, the compliance of the original data sources with the Findable, Accessible, Interoperable, Reusable (FAIR) data principles as well as the ability to semantically map feature names to common vocabularies such as MedGen, PubChem, and Ensembl gene IDs was essential (*14*, *15*). We are aware of a certain degree of redundancy among the data dimensions as for example pathway associations are derived from gene associations. Nonetheless, pathways and genes represent different levels of organization and thus of information as two diseases that share associated pathways can be associated with different genes. Also, we demonstrate that the pathway dimension generates a different disease similarity map as compared to the gene dimension, although both show relatively high congruence. We name entities and the links between them uniformly. Genes and proteins are summarized under the entity of genes. Both approved drugs and non-drugs are summarized under the entity of chemicals, which also includes biologicals such as therapeutic antibodies. With the chemical-medicating-disease links we cover the dimension of approved drugs and therefore beneficial links between chemicals and diseases. The other two chemical dimensions are largely undirected and might represent (as well) potentially negative effects such as side effects of drugs and disease-triggering effects of chemical substances.

### 2.1 Mortality drives the frequency of disease-associated features

We hypothesize that features associated with a high number of diseases *i.e.*, very common disease features, might be relevant to discriminate broad etiological groups of diseases such as cancerous versus non-cancerous diseases, while less common features help to refine the discrimination of similar diseases. To obtain an overview of the commonness of disease features, we counted how often a gene, pathway, symptom, and chemical is linked to diseases and ranked them according to the frequency (Figure 2, Table S1).

**Figure 2:**
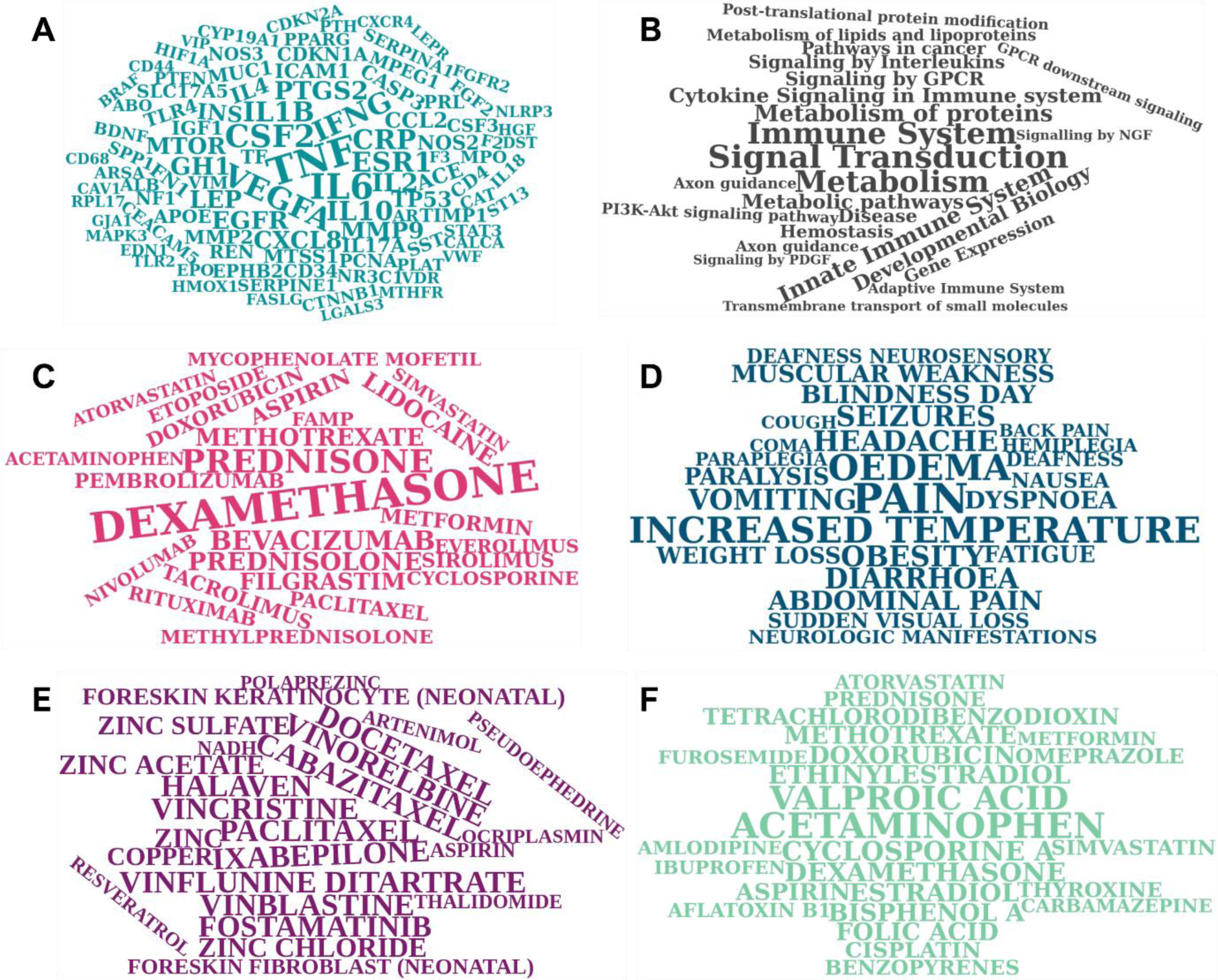
Frequencies of features being linked to diseases. For each of the six data dimensions, we counted how often single features (*i.e.*, genes, pathways, chemicals) are linked to diseases as shown in Figure 1. The wordclouds show the top 100 frequent features for each dimension. **A.** The 100 genes that are most frequently associated with diseases. *TNF* is associated with the highest number of diseases (2,490 of 7,618, 33%). **B.** The 25 pathways that are most frequently associated with diseases. The top pathway, Signal Transduction, is associated with 1,887 of 4,894 diseases (39%). **C**. The 25 chemicals that are most frequently used to treat diseases. The top chemical Dexamethasone is associated with 242 of 1811 diseases (13%). **D.** The 25 symptoms most frequently associated with diseases. The top symptom, pain, is associated with 2,290 of 4,579 diseases (50%). **E.** The top 25 chemicals targeting genes associated with diseases. The top chemical Paclitaxel is linked to 5,169 of 7188 diseases (72%). **F.** The 25 chemicals most frequently associated with diseases. The top chemical Acetaminophen is associated with 7,967 of 16,723 diseases (48%).

The genes most often associated with diseases are *TNF*, *IL6*, *CSF2*, *VEGFA*, and *IFNG.* They are pivotal genes in immune functions, which explains why they are associated with a high number of diseases as immune functions present an overarching system of both physiological and pathological cellular processes. The drugs that are medicating the highest number of diseases are Dexamethasone, Prednisone, Bevacizumab, Prednisolone, and Methotrexate, which is in accordance with the gene level results, since these drugs primarily target immune pathways. Besides the three corticosteroids which are used as first-line medication for multiple illnesses, we retrieved Bevacizumab, a monoclonal antibody targeting VEGF and approved for various cancers, as well as Methotrexate, which is approved for multiple cancerous and rheumatoid diseases. The chemicals associated with the highest number of diseases are Acetaminophen (paracetamol), Valproic acid, Cyclosporine A, Dexamethasone, and Ethinylestradiol. These chemicals are frequently used pharmaceuticals. Thus, they might be associated with a high number of diseases due to (1) multiple known or suspected side effects and/or (2) due to their apparently broad applicability. The chemicals linked to the highest number of diseases via targeting disease-associated genes are Paclitaxel, Cabazitaxel, Ixabepilone, Vincristine, and Vinflunine. All of them are cytostatics. Thus, they present impactful perturbations affecting a high number of proteins. For example, Paclitaxel is targeting about 1,500 genes such as *TNF* and tubulins, which are on the other hand associated with various diseases. In summary, features associated with a high proportion of diseases might only be helpful to map diseases at a broader level, whereas not for local fine-tune mappings.

We hypothesize that the number of diseases a gene, symptom, or chemical is related to is influenced by the extent of research globally performed on it. To investigate this relationship, we retrieved curated data from NCBI about how often a gene, a chemical, or a disease is linked to a publication in PubMed. We investigated the relationship between the number of publications and the number of disease associations. We observed a moderate correlation between the number of publications and the number of disease associations for genes (R=0.7) and symptoms (R=0.65), respectively. This suggests, not surprisingly, that genes and symptoms associated with many diseases tend to be mentioned in publications more frequently (Figure 3 A, B). The genes *TP53* and *TNF* present two opposing extremes. *TNF* has the highest number of disease associations, likely given its key role in inflammation and response to injury. However, it is surprisingly moderately mentioned in publications, likely because the research is dispensed on multiple TNF-family members. *TP53* has by far the highest number of related publications, but only a moderate number of disease associations. While *TP53* is almost exclusively associated with cancerous diseases, cancers account for the majority of biomedical publications among the disease categories (*16*). Further, *TGFB1* and *APOE* show a remarkably high number of publications despite moderate numbers of disease associations. A possible explanation is their association with yet incurable and fatal conditions such as organ fibrosis and Alzheimer disease (*17*, *18*). Two symptoms, namely pain and obesity, show an outstanding number of publications, with pain also being the symptom associated with the highest number of diseases. Several symptoms deviate from the overall correlation trend between disease associations and publications. Neurological manifestations, for instance, are associated with a moderate number of diseases, but have a notably high number of publications, likely due to their potential severity as compared to vomiting, which is associated with a high number of diseases but less mentioned in publications. Noteworthy, while neurological manifestations comprise multiple symptoms, vomiting is a precise definition of a symptom. Thus, semantic qualities can particularly bias clinical features of diseases. For chemicals, we observed a weak correlation between the number of disease associations and the number of publications suggesting that our understanding of human diseases caused by specific chemical exposure is still broadly incomplete (Figure 3 C). Notably, the chemicals considered in this study consist of a broad and heterogenous group of molecules in contrast to genes. Likewise, associations between chemicals and diseases are of various types that go beyond “is harmful” and “is medicating”. The chemicals associated with the highest number of diseases such as Acetaminophen and Valproic acid are used to medicate diseases, thus by design, they are impactful. In contrast, such substances mentioned in publications most frequently (oxygen, glucose, and water) elude from the correlation between disease associations and publications due to their harmlessness.

**Figure 3:**
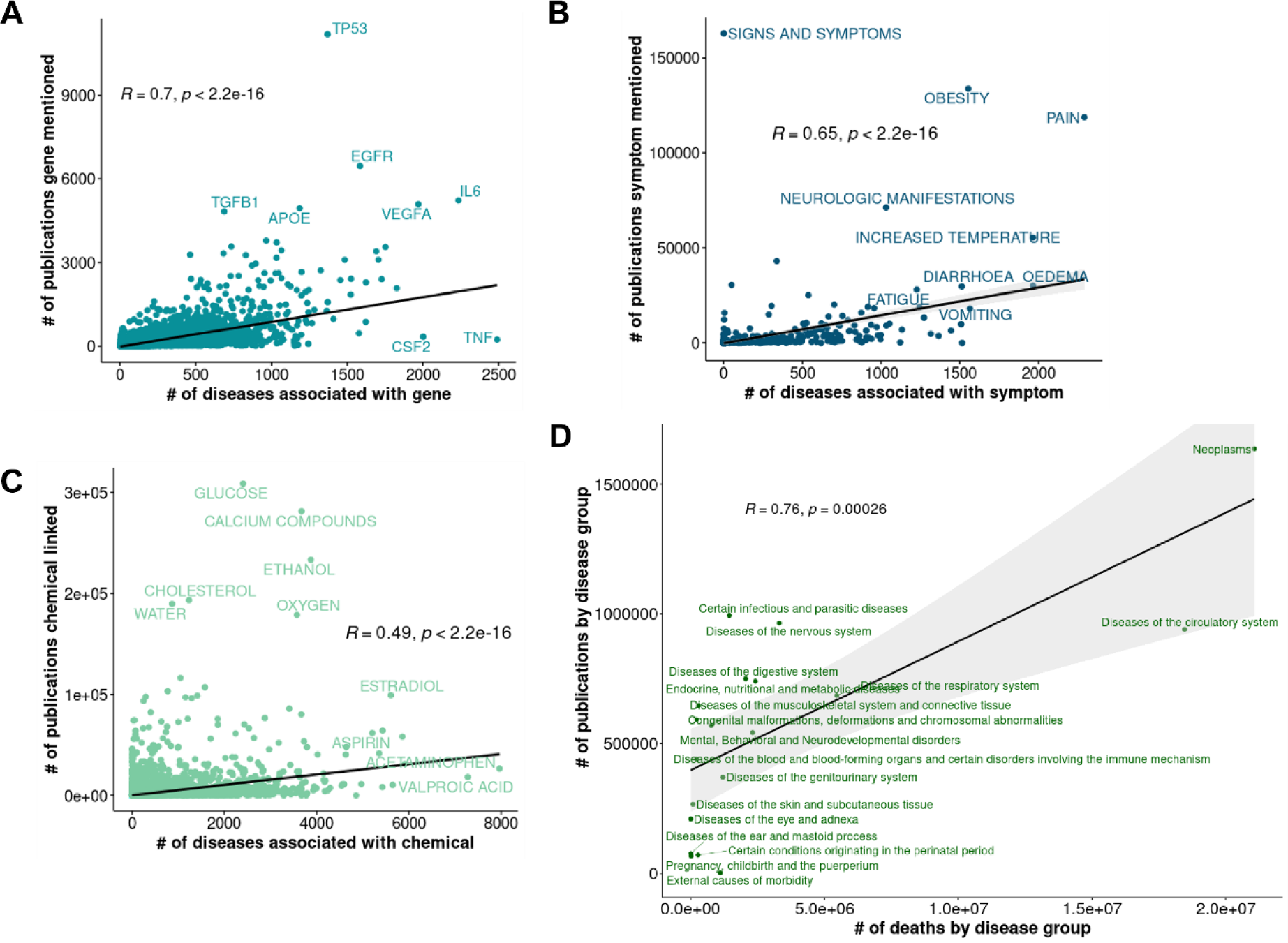
Relationship of disease associations and publications. Data of how often a gene, chemical or disease is linked to a publication was retrieved from NCBI. **A.** The number of publications for a gene (y-axis) was compared with the number of diseases associated with the gene (x-axis). **B.** The number of publications for a symptom (y-axis) was compared with the number of diseases associated with the symptom (x-axis). **C.** The number of publications for a chemical (y-axis) was compared with the number of diseases associated with the chemical (x-axis). **D.** Diseases were grouped according to ICD-10 chapters. The number of publications for a disease group (y-axis) was compared with the number of deaths caused by the disease group as recorded in the United States between 1999 and 2020 (x-axis). Spearman correlation coefficients and p-values are indicated in the plot areas. # = number.

Based on the observed correlation between the number of publications and the number of disease associations, a potential literature bias especially for gene-disease associations needs to be considered. To infer the causality in the observed correlation, we retrieved mortality data for diseases from the WHO and investigated the relationship between mortality and number of publications for groups of diseases as defined by ICD-10 chapters. The overall mortality of diseases (in the United States between 1999 and 2020) correlates with the number of publications for diseases (Figure 3 D). This suggests that the number of deaths caused by a disease, *i.e.*, its overall burden, drives the extent of research performed on a disease and thus the number of publications. Altogether, these observations suggest that mortality drives the extent of research performed on disease-associated features such as genes and chemicals.

### 2.2 Disease relationships based on six data dimensions

Next, we aimed to explore disease relationships based on the associated features. To compare the six data dimensions, we focused on a subset of diseases present in all data layers. The number of diseases with available data varied largely among the six data dimensions. In each dimension, we first excluded diseases with a number of associated features falling in the lower quartile of the distribution. After this filtering, the chemical dimension comprised 12,666 diseases, while for the dimension of the symptoms, only 3,587 diseases were available (Figure 4 A). To compute disease-to-disease relationships, we used 502 diseases available in each dimension.

**Figure 4:**
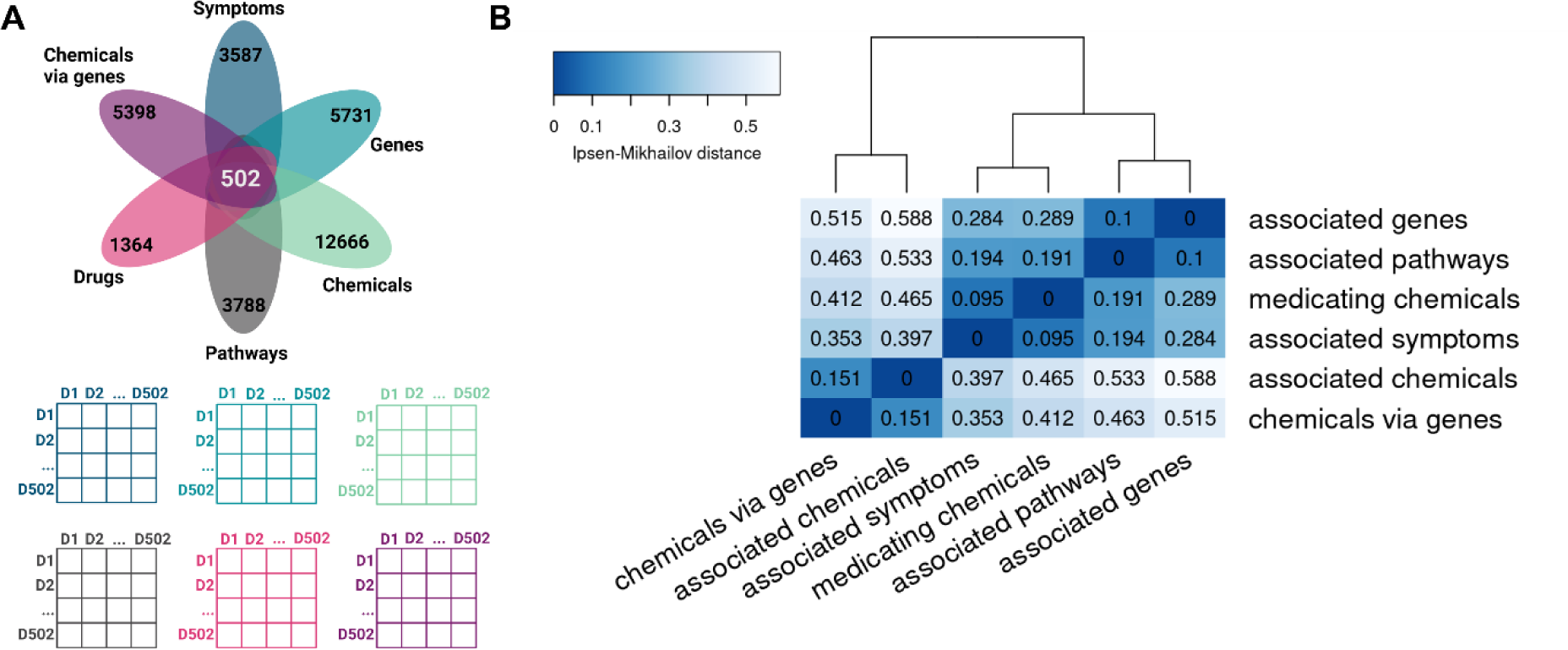
Disease-to-disease distance matrices in six data dimensions. **A.** Top: The number of diseases with available data for each dimension and the overlap among them. Prior to the depicted numbers, diseases with a low number of associated features (lower quartile) had been removed from each dimension. Bottom: For each dimension, a distance matrix was computed based on a consensus of six distance measures. **B.** Hierarchical clustering of the distance matrices in the six dimensions based on the Ipsen-Mikhailov distance computed among the matrices.

We employed a consensus approach utilizing six distinct distance measures to calculate pairwise distances between diseases within each data layer. This resulted in the creation of a separate distance matrix for each dimension (Supporting Data 2). To compare the six distance matrices, we computed the Ipsen-Mikhailov distance among them (Figure 4 B).

The 502 diseases cluster similarly based on the gene and pathway dimension, mirroring the fact that associated pathways represent functionally homogeneous sets of disease-associated genes. Likewise, the clustering based on the associated symptoms resembles the one based on medicating drugs, indicating a close relationship of the clinical dimensions among each other. Eventually, drugs are treating a certain group of symptoms, the combinations of which have been labeled as a distinct disease. In the two dimensions of chemicals, the 502 diseases are overall clustered differently as compared to the other four dimensions, as the Ipsen-Mikhailov distance between those and the clinical, gene and pathway dimensions is around 0.5. Since no data layer completely resembles another, we conclude that each layer contributes unique information to a certain extent.

The ICD-10 classification system groups diseases based on the affected anatomical locations and broad etiological groups. To investigate if the ICD-10 classification mirrors disease relationship beyond anatomy, we compared the patterns of disease relatedness in the ICD-10 with those emerging from the six data dimensions curated here. We observed that the clustering based on the ICD-10 codes yields a broadly different disease map as compared to all the other dimensions as well as the consensus of the six dimensions (Figure S1, Supporting Data 2). This indicates that the ICD-10 classification does not completely cover the information of disease relationships gained from any of our chosen dimensions. In conclusion, diseases cluster very differently on the mechanistic level as compared to a largely anatomical level. Hence, the ICD-10 classification of diseases is practical for clinical and regulatory affairs since it is an intuitive system that reflects broad phenotypic criteria. However, according to our observations, it largely ignores aspects regarding the mechanism of action of diseases.

Interestingly, our findings do not replicate the observations made by Sakaue *et al*., whose atlas of genetic associations supports the ICD-10 classification (*19*). However, in that study, only genetic associations as measured by Genome-wide association studies (GWAS) were considered. Here, instead, we considered multiple dimensions including but not limited to genetic associations, which extends the potential etiological knowledge to pathogenetic pharmacological and clinical ones. Notably, also our dimension of associated genes comprising information gathered from GWAS (among others) does not coincide with the ICD-10 classification. In another study, Zitnik *et al*. reported overlaps between systems-level molecular layers, such as genes, drugs, and protein-protein interactions, with existing disease classifications that however partially include characteristics that go beyond anatomical locations (*10*). Here instead, with the ICD-10, we focused on a system at the far most anatomical pole.

### 2.3 Type 2 diabetes and Alzheimer disease share rarely associated features

We investigated the disease landscape of three common diseases in the Western population such as psoriasis, asthma, and type 2 diabetes (T2D) in more detail looking at their immediate neighbors according to the computed consensus distance. All three diseases have inflammation among their top ten similar diseases indicating that they share this as a central characteristic (Figure 5). Further, psoriasis and asthma share rheumatoid arthritis and allergy among their top similar diseases, while asthma and T2D share hypertension and myocardial infarction.

**Figure 5:**
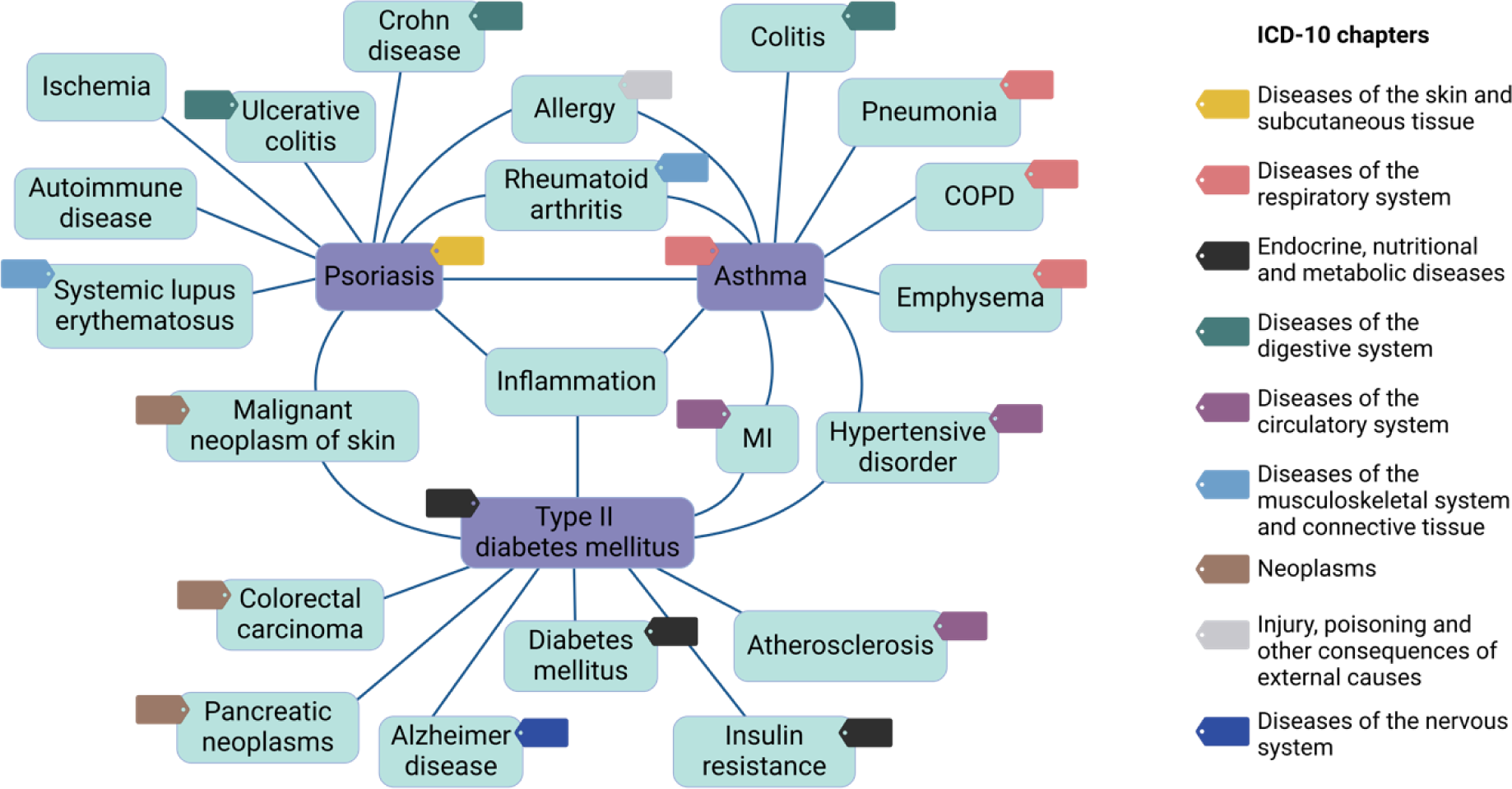
Common diseases and their top ten similar diseases. Three common diseases in the Western population (violet boxes) and their top ten similar diseases (turquoise boxes) according to the compiled consensus distance score of six data dimensions. Edges indicate if a disease is among the top ten similar diseases. Colored tags indicate the ICD-10 chapter the disease belongs to. MI: Myocardial infarction; COPD: Chronic obstructive pulmonary disease

The ten most similar diseases to psoriasis are autoimmune disease, allergy, malignant neoplasm of skin, inflammation, rheumatoid arthritis, Crohn disease, ulcerative colitis, ischemia, asthma, and systemic lupus erythematosus. This spectrum covers four distinct branches of the ICD-10 system (without considering the generic disease terms “autoimmune disease”, “inflammation”, and “ischemia”) further highlighting the gap between anatomical and mechanistic disease maps (Figure 5, Table S2). In the ICD system, psoriasis is assigned to the group “Diseases of the skin and subcutaneous tissue”. Thus, it is noteworthy that the immediate neighborhood of psoriasis is characterized by (auto)-inflammatory diseases, whereas only one neoplastic skin disease is among the top ten similar diseases of psoriasis. However, the high overall similarity between psoriasis and malignant neoplasms of the skin can be explained by hyperproliferation of keratinocytes as a shared fundamental mechanism of action (*20*).

The ten most similar diseases to asthma according to the computed consensus distance are allergy, chronic obstructive pulmonary disease (COPD), inflammation, pneumonia, myocardial infarction, colitis, rheumatoid arthritis, psoriasis, emphysema, and hypertension. Those diseases cover five disease groups in the ICD system (Figure 5, Table S2). We observed multiple diseases of the respiratory system among the closest diseases to asthma indicating that asthma is mechanistically closer to other lung diseases, while this does not seem to apply for psoriasis with respect to skin diseases supporting the paradigm of psoriasis as a systemic disorder (*21*).

The ten most similar diseases to T2D according to the consensus distance are diabetes mellitus, hypertension, colorectal carcinoma, insulin resistance, myocardial infarction, inflammation, Alzheimer disease, malignant neoplasm of skin, pancreatic neoplasms, and atherosclerosis. While diabetes mellitus and insulin resistance present related diseases or precursors, the high similarity of T2D with three cancerous diseases as well as cardiovascular diseases is in accordance with epidemiological observations. Individuals with T2D have a higher risk of developing colorectal cancer as well as endpoints of atherosclerosis such as stroke and myocardial infarction (*22*). However, whether T2D can be considered an exposure variable causing a higher lifetime risk of colorectal cancer as well as atherosclerosis or whether the epidemiological association is due to shared risk factors such as obesity and lifestyle factors is not yet completely understood (*23*, *24*). Nonetheless, we find it remarkable that our disease relationship model reflects epidemiological associations. Further, the observed closeness of T2D and Alzheimer disease is intriguing and indicates the power of multi-dimensional but non-anatomical disease views. There is increasing evidence of an association between T2D and Alzheimer disease (*25*). An altered glucose metabolism and insulin signaling are possible factors linking the two diseases, but mechanistically, the link is poorly understood. To investigate the mechanistic link between them, we extracted features shared by those two diseases. To isolate shared features from common inflammatory features, we excluded features that are associated with psoriasis, asthma, and/or inflammation. The 73 associated pathways shared by Alzheimer disease and T2D, but not psoriasis, asthma and/or inflammation include pathways such as regulation of insulin secretion, insulin receptor recycling, insulin processing, glucose transport, glycolysis, zinc transporters, zinc efflux and compartmentalization, glycerophospholipid metabolism, and Notch signaling. Notably, the pathway “Zinc efflux and compartmentalization by the SLC30 family” is associated only with three further diseases (Jacksonian seizure, Esophageal neoplasm, and transient neonatal zinc deficiency) of all 4,894 diseases available on the pathway data layer. The pathway “Zinc transporters” is associated with 14 further diseases underlining that T2D and Alzheimer share some overall extremely rare features that might assist in further elucidating their mechanistic link.

### 2.4 Diseases cluster into functionally and mechanistically coherent groups

Next, we aimed at exploring groups of similar diseases and the features that discriminate each disease group from the others. Therefore, hierarchical clustering of 502 diseases was conducted based on the consensus distance score calculated from pairwise disease-to-disease distances in six data dimensions. To identify the optimal number of disease clusters, the Dunn index was computed which yields higher values for low intra-cluster and high inter-cluster distances. Overall and expectedly, the Dunn index is increasing with increasing the number of clusters, while 71 to 78 clusters reached a local maximum (Figure 6, A). We proceeded with 71 clusters of diseases and tested by Fisher’s exact test the enrichment of features within disease clusters.

**Figure 6:**
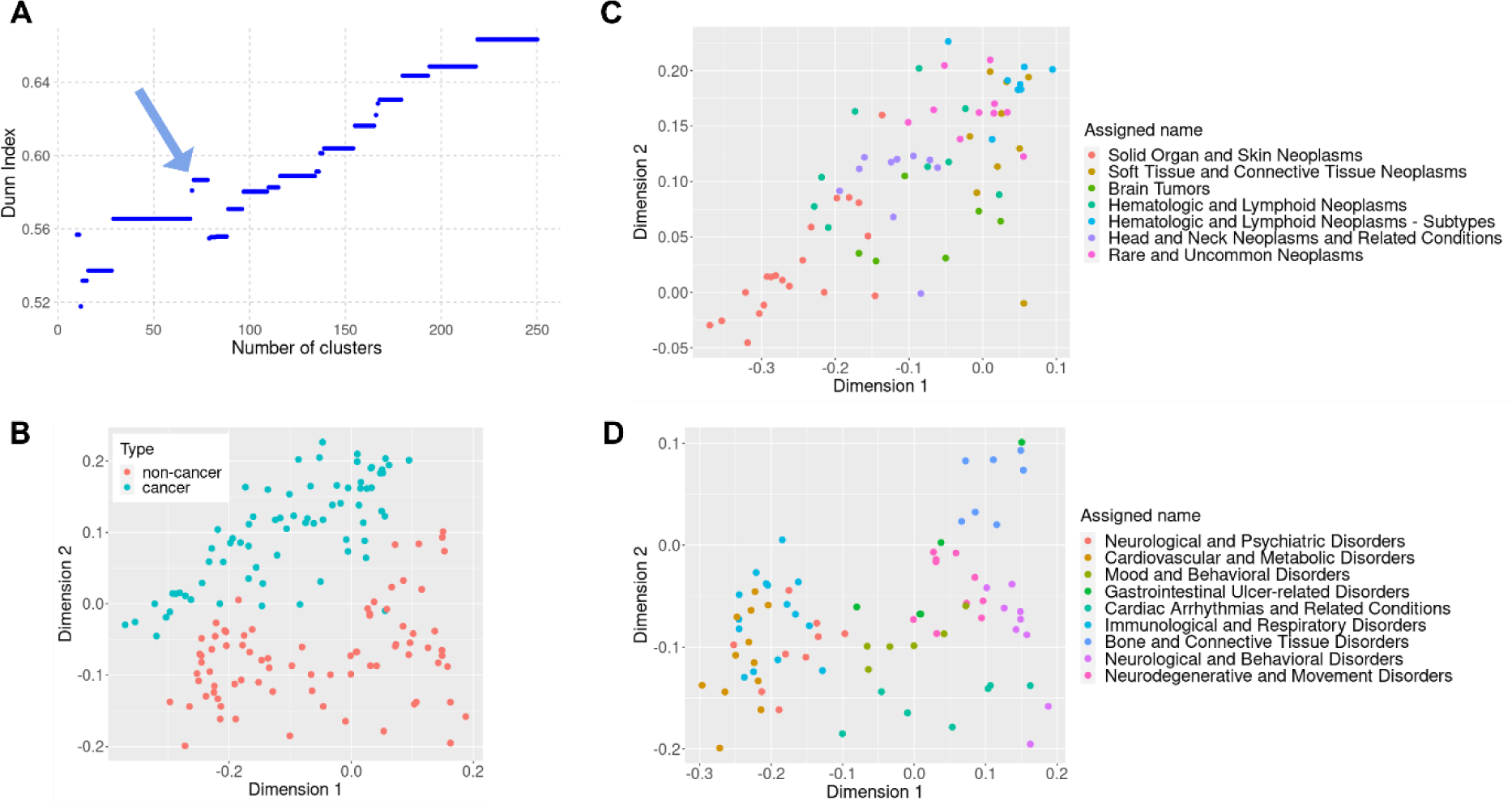
Disease clusters based on consensus distance. **A.** After hierarchical clustering of 502 diseases based on the consensus distance score calculated from six data dimensions, the tree was cut into 2 to 250 clusters and the Dunn index was computed for the different groupings. There was a local maximum for 71 to 78 clusters (blue arrow), and we therefore proceeded with our analysis using 71 disease clusters. **B.** Among 16 candidate clusters of interest, seven clusters comprised cancerous diseases (blue) and nine clusters comprised non-cancerous diseases (red). Multidimensional scaling (MDS) separated the two types of clusters. **C.** The seven cancerous disease clusters in an MDS plot with cluster names as suggested by ChatGPT. **D.** The nine non-cancerous disease clusters in an MDS plot with cluster names as suggested by ChatGPT.

#### Cancerous and non-cancerous disease clusters are two distinct “continents” on the multi-dimensional disease map

We let ChatGPT assign an overall best describing name for each of the 71 disease clusters (Table S2, Figure 6 C, D). Hereafter, we focused on 16 candidate clusters containing at least five diseases that represented a homogenous mix of specific diseases, while the remaining 55 clusters were small and/or contained a heterogenous mix of symptoms and generic phenotypes. Seven of the 16 candidate disease clusters comprised almost exclusively neoplasms (both benign and malignant), while the other nine candidate clusters exclusively comprised non-cancerous diseases. Representation in two dimensions by multidimensional scaling separates those two types of clusters completely indicating the presence of characteristics frequently associated with cancer, while not or rarely being associated with non-cancerous diseases (Figure 6, B). The top five enriched genes in the seven cancer clusters as compared to the nine non-cancer clusters were *BCOR* (BCL6 corepressor), *TOP2A* (DNA topoisomerase II alpha), *MDM2* (MDM2 proto-oncogene), *CCND1* (cyclin D1), and *POLD1* (DNA polymerase delta 1, catalytic subunit). Among in total 161 over-represented genes in the cancer clusters were as well the usual suspects *TP53*, *KRAS*, *BRAF*, and *MYC*, and genes encoding the beta tubulin protein domains (Supporting Data 4). In contrast, the 27 over-represented genes in the non-cancer clusters as compared to the cancer clusters are involved in Neuroactive ligand-receptor interaction and glutamate receptor pathways. Interestingly, genes of these pathways were not only associated with classical nervous system diseases as one could assume, but also with various metabolic and inflammatory diseases such as diabetes, obesity, hypertension, ischemic disorders, and inflammatory bowel diseases indicating that the enriched genes in the non-cancer clusters are not exclusively driven by the (high) fraction of nervous system-related disorders. While in the cancer clusters, no symptoms were significantly over-represented, in the non-cancer clusters, nervous system-related symptoms such as dyskinesia and tremor were over-represented, but also less specific symptoms such as sleep deprivation and increased weight. Overall, our observations about differences between cancerous and non-cancerous disease clusters are not novel, however, they might question existing gene popularity in cancerous and non-cancerous diseases.

#### Cancer disease clusters are driven by cellular and tissue origin

Interestingly, in most of the seven indicated cancer clusters, there exists one clearly predominating type of cellular origin namely either epithelia, connective tissues, blood forming tissues, or nervous tissues. The two most contrasting cancer clusters were a cluster of mainly sarcomas named “Soft Tissue and Connective Tissue Neoplasms” and a cluster of mainly carcinomas named “Solid Organ and Skin Neoplasms” (Table S2). In multidimensional-scaling two-dimensional space, those two clusters are completely discrete suggesting that carcinomas and soft tissue cancers represent two poles of neoplasms (Figure 6, C). Two clusters of hematologic and lymphoid neoplasms represented cancers from the blood forming tissues, with one of them clustering with the soft tissue cancers, while the other was less homogeneous and overlapped with the cluster of solid tumors. Notably, a cluster named “Rare and Uncommon Neoplasms” represented a relatively dense cluster in two-dimensional space, which was located towards the soft-tissue cancer pole, while the cluster named “Head and Neck Neoplasms and Related Conditions” extended between the two poles of carcinomas and soft tissue cancers. The small cluster of neoplasms evolving from nervous tissues (“Brain tumors”) formed a heterogeneous group of diseases between the two poles, as well.

In a more specific context, the cluster named “Soft Tissue and Connective Tissue Neoplasms” comprised multiple sarcomas, fibroma presenting a benign precursor to a sarcoma, as well as Schwannoma indicating that their cellular origin dictates their similarity rather than anatomical locations. One gene, *MYOG* (myogenin), was significantly enriched in this disease cluster as compared to the background of diseases not part of the cluster (Supporting Data 3). This transcription factor involved in differentiation of mesenchymal cells into skeletal muscle has long been known as marker for rhabdomyosarcoma, however it has been associated with other types of sarcomas, as well (*26*, *27*). Facial paralysis was the strongest enriched symptom in this disease cluster (non-significant after correction for multiple testing). Interestingly, this symptom was not enriched in clusters with multiple enriched neurological symptoms such as the cluster of brain tumors, neurodegenerative and movement disorders, and neurological and behavioral disorders. The drug Voruciclib was the strongest enriched substance among those that target genes associated with this cluster of soft tissue neoplasms. While Voruciclib is currently tested as treatment for hematologic malignancies (NCT03547115), it was not enriched in clusters of such. Our results indicate that the targets of Voruciclib have implications particularly in the pathology of soft tissue cancers suggesting a potential use for the treatment of solid tumors beside hematologic cancers.

In contrast to the cluster of cancers derived from soft tissues, the cluster named “Solid Organ and Skin Neoplasms” comprised cancers predominantly derived from epithelial linings of various organs such as lungs, stomach, pancreas, liver, colon, uterus, prostate, and ovaries. In this disease cluster, osteosarcoma and melanoma present two non-carcinomas, while the latter is still closely associated with epithelial tissues (Table S2). Beside endometrial neoplasms, endometriosis as the only non-cancerous disease fell into this cluster indicating a strong relatedness to cancerous diseases especially carcinomas. In total, 1,431 genes and 714 pathways were enriched in this cluster as compared to the diseases not part of this cluster showcasing the strong molecular fingerprint of carcinomas as a highly distinct disease group (Supporting Data 3). The top enriched genes are overall involved in the cell cycle machinery. Enriched symptoms in this cluster included hemoptysis, intractable pain, and feminization.

While the cluster “Head and Neck Neoplasms and Related Conditions” indeed showed some accumulation of this anatomical site, with diseases mostly evolving from epithelia, a striking characteristic of this cluster was the accumulation of precancerous conditions (oral submucous fibrosis, precancerous condition), conditions that pave the way to cancer (gastroesophageal reflux disease), and benign neoplasms (papilloma) as well as cancers that rarely metastasize, grow slowly or are often diagnosed at early stages (basal cell carcinoma, neoplasm of the thyroid gland) (Table S2) (*28*, *29*). The factor that diseases in this cluster are overall more likely to be detected as they are visible (mouth neoplasms, neoplasm of the tongue) or symptomatic early (thyroid cancer, reflux), might contribute to the shared characteristics of diseases in this cluster or even be the key determinator. Noteworthy, one disease in this cluster clearly not matching any of the mentioned criteria (precancerous, benign, early detection) is Cholangiocarcinoma. On the gene and symptom level, there were no enriched features in this cluster indicating a certain level of unspecificity and/or heterogeneity. On the pathway level, signaling by FGFR was the striking characteristic of this cluster, likely related to the implication of profibrotic processes in the development and progression of those cancerous and pre-cancerous diseases. Among the strongest enriched chemicals being associated with diseases in this cluster are substances such as Benzidine and Hydrazobenzene, which are classified as certainly oncogenic.

Altogether, our observations suggest that neoplasms group according to their cellular compartment of origin, with notable differences between a group of carcinomas and a group of sarcomas.

#### A link between cardiometabolic-inflammatory and nervous system related conditions

Among the nine candidate clusters exclusively comprising non-cancerous diseases, three (larger) clusters named “Neurological and Psychiatric Disorders” (n=9 diseases), “Cardiovascular and Metabolic Disorders” (n=12), and “Immunological and Respiratory Disorders” (n=15) largely overlapped and formed a neuro-metabolic-inflammatory supercluster in two-dimensional MDS plot (Figure 6, D, Table S2). Completely discrete from this were the clusters named “Neurological and Behavioral Disorders” (n=9) and “Neurodegenerative and Movement Disorders” (n=10). At the interface between those two neuro-clusters and the neuro-metabolic-inflammatory supercluster was a group of diseases named “Mood and Behavioral Disorders” (n=6) comprising different forms of depressive disorders and attention deficit hyperactivity disorder (ADHD). The clusters of “Cardiac Arrhythmias and Related Conditions” (n=7) and “Bone and Connective Tissue Disorders” (n=7) represented two distinct clusters that showed little overlap with other clusters indicating a certain level of organ- and tissue-specificity for those disease groups. Interestingly, the diseases of the cluster “Gastrointestinal Ulcer-related Disorders” (n=5), which in classical disease systems, would be grouped closely together due to their phenotypic similarity, spread relatively broadly based on the calculated consensus distance.

The closeness of the clusters “Cardiovascular and Metabolic Disorders” and “Immunological and Respiratory Disorders” supports more recent epidemiological findings and definitions of metabolic-chronic-inflammatory risk profiles. The link between inflammatory and immune system related disorders such as psoriasis and rheumatoid arthritis, and cardiometabolic diseases such as atherosclerosis and diabetes is hypothesized to result from shared risk factors and/or causal relationships that favor certain disease trajectories (*30–32*). In both scenarios, mechanistic similarities between immune system related and cardiometabolic diseases seem to be reasonable. Remarkably, the cluster “Neurological and Psychiatric Disorders” containing the diseases Amyotrophic lateral sclerosis, late-onset Parkinson disease, pain, Alzheimer disease, Ischemic Encephalopathy, Nicotine dependence, Schizophrenia, Jacksonian seizure, and Status epilepticus has overlapping characteristics with this metabolic-chronic-inflammatory disease group according to our results. We therefore investigated in more detail what those three clusters have in common that separates them from the diseases outside of those three clusters as well as what are differences among those three clusters. The three clusters shared 53 associated genes that were enriched in each of them as compared to the diseases that did not fall in the specific cluster (Supporting Data 3). These genes, which included seven genes encoding GABA(A) receptors, *TACR1* encoding the tachykinin receptor 1 and its ligand *TAC1*, as well as *NTS* (neurotensin), revealed the strongest enrichment for a gene set summarized as neuroactive ligand-receptor interaction. Further, the union of the enriched genes of the three clusters strongly accumulated in this gene set as well, suggesting that neuroactive signaling might be a link between neuro-metabolic, cardio-metabolic and immune-related malfunctioning (Figure S2). To identify differentiating features between the three clusters, we performed a direct pairwise comparison (Supporting Data 5). In the cluster of “Immunological and Respiratory Disorders” the genes *IL12RB2*, *CIITA* (class II major histocompatibility complex transactivator), *TNFSF4*, *MMP19*, *TIGIT* (T cell immunoreceptor with Ig and ITIM domains), *KLRC1* (killer cell lectin like receptor C1), and *IL17C* were overrepresented (nominal p-value <0.05) as compared to the “Neurological and Psychiatric Disorders”. In the cluster of “Cardiovascular and Metabolic Disorders” the genes *TIMM8A (*translocase of inner mitochondrial membrane 8A), *RBPJ* (recombination signal binding protein for immunoglobulin kappa J region), *BDKRB2* (bradykinin receptor B2), *NPR2* (natriuretic peptide receptor 2), *PLN* (phospholamban), *FABP1* (fatty acid binding protein 1), and *NOD1* (nucleotide binding oligomerization domain containing 1) were overrepresented (nominal p-value <0.05) as compared to the “Neurological and Psychiatric Disorders”. Genes such as *IL12RB2*, *IL17C*, *KLRC1, NOD1*, and *TNFSF4* are involved in the interplay between innate and adaptive immune responses including defense at epithelial surfaces, recognition of self and antigens, as well as antigen presentation, indicating that these processes partially involve different key players in immunologically privileged organs like the brain. However, cells of the central and peripheral nervous system share multiple gene signatures and functions with other organs such as the liver and gastrointestinal tract as shown by recent single cell atlases, and this might explain shared a mechanistic basis of metabolic-inflammatory and classical neurological disorders (*33*, *34*).

Overall, we observed that several disease clusters are characterized by a predominating ICD-10 super-concept such as the cancer clusters and non-cancer clusters such as “Cardiac Arrhythmias and Related Conditions” and “Bone and Connective Tissue Disorders”. However, clusters like the “Immunological and Respiratory Disorders” and “Cardiovascular and Metabolic Disorders” break classical phenotypic and anatomy-driven disease classes, as they comprise multiple ICD-10 chapters. Furthermore, the closeness of the latter two with a cluster of neurological and psychiatric disorders supports a novel view of disease categories. We believe that approaches like ours offer new perspectives on disease etiologies and the potential of reformulating therapeutic interventions or considerations of new treatment strategies. A revision of current treatment strategies is needed to address the rising challenge to treat multimorbidity. In addition, the drug target space needs to be expanded as it was recently shown that current clinical trials focus on previously tested drug targets and cover only a fraction of druggable proteins (*35*). Multi-dimensional maps of diseases can assist in solving both problems. First, a multi-layer data-driven comparison of diseases can identify new disease relationships and thus possibilities of shared treatment options, and second, identified disease groups and characteristics like associated genes that are enriched in such, can reveal novel and specific biomarkers that might serve as drug targets for drug development. Further, we emphasize that approaches like ours can be shifted from the epidemiological towards an individual patient level whenever patient data are available on a larger scale. In this way, multimorbid patients could be grouped completely disease-agnostic according to their clinical and molecular variables, and further, endotypes of a specific disease can be leveraged by systematically comparing patients on multiple dimensions.

## 3 Conclusions

By mapping diseases based on multiple data dimensions we gain a holistic view of disease landscapes, allowing us to identify new families of diseases as well as key features that suggest shared mechanisms of action. We conclude that the applied methodology is overall robust, as it captured several patterns that are in accordance with current biomedical knowledge such as the distinct grouping of cancer and non-cancer clusters and the similarities of metabolic and inflammatory conditions. While our approach is limited by the absence of performance assessment strategies, we refer to the exploratory nature of those kinds of studies with overall missing reference systems. A further constraint of our study is the existence of non-unified terminologies of disease names limiting the number of diseases and dimensions that can be studied. We observed striking differences between the here constructed disease map and the ICD-10 classification of diseases, albeit some identified disease groups correlate with ICD-10 super-concepts such as clusters of exclusively cancerous diseases and clusters of diseases mainly manifesting as conditions of the nervous system. Our results highlight limitations of traditional anatomical and broad etiological classifications and emphasize the potential of functional and mechanistic relationships of diseases to update existing classifications or as a parallel disease universe. Systematic mapping of diseases considering individual disease features of various dimensions provides a solution to fill the gap between epidemiological and personalized models, which is needed to address medical challenges such as the rising prevalence of multimorbidity and low drug efficacy rates due to currently applied pharmacology approaches. Hence, a revision of established disease views might entail implications for drug discovery processes as well as it might assist in elucidating patterns of comorbidities and in reformulating existing treatment approaches. In summary, a data-driven update of existing disease classifications presents promising avenues for future research and clinical trial design to overcome largely phenotypic and stiff disease views.

## 4 Experimental Section

### Data sources and retrieval

We used publicly available data linked to diseases of six different dimensions: (1) associated genes were retrieved from Open targets (retrieved 15/02/2020) (*36*), (2) associated pathways were retrieved from KEGG (retrieved 14/10/2021) and the Comparative Toxicogenomics Database (CTD) (retrieved 04/10/2019) (*37*, *38*), (3) associated symptoms were retrieved from a study by Zhou *et al.* (*39*), (4) drug profiles were retrieved from Open targets (retrieved 15/02/2020), LabeledIn (retrieved 11/02/2020), and a study from Wang *et al.* (*36*, *40*, *41*), (5) associated chemicals were retrieved from CTD, SIDER, offsides and MEDI (retrieved 04/10/2019, 07/07/2020, 25/06/2020, and 02/07/2020, respectively) (*38*, *42–44*), (6) chemicals targeting disease-associated genes represented the intersection of chemical-target information retrieved from DrukBank (retrieved 11/06/2021), Open Targets (retrieved 15/02/2020), and Pharos (retrieved 05/11/2020) and associated genes retrieved from Open targets (retrieved 15/02/2020) (*36*, *45*, *46*). The data were retrieved from a knowledge space recently published in which publicly available data has been reposited in a graph-based dataspace after extensive curation and homogenization (*13*). For the integration of various data sources into this graph-based database structure, unified vocabularies were used such as NCBI PubChem identifiers for chemicals, Ensembl gene identifiers for genes and their products, and NCBI MedGen concept identifiers for diseases. The disease codes of the ICD-10 system were retrieved from the ftp server of the Centers for Disease Control and Prevention (CDC). To match ICD-10 codes with NCBI MedGen concept identifiers, we used the NIH Unified Medical Language System (UMLS) Terminology Services (*5*). The UMLS obtained 5,141 matches of ICD-10 codes and MedGen concept identifiers. However, this included matched pairs such as C0005604:P10-P159 and C0004096:J45, which are non-exact matches. In both cases, a MedGen concept ID was not assigned to one exact ICD-10 code, but to a superordinate ICD-10 group. Furthermore, for multiple MedGen IDs, there are no matches at all to the ICD-10 system. Due to this, the comparison of the six data dimensions with the ICD-10 system was based on only 131 of the initial 502 diseases. Summary data for MedGen and PubMed identifiers (retrieved 13/04/2023), chemical identifiers and PubMed identifiers (retrieved 18/04/2023), and Gene identifiers and PubMed identifiers (retrieved 14/02/2023) were retrieved from the ftp server of NCBI (*15*). Mortality data were retrieved from the Mortality database of the WHO (retrieved 03/10/23). For analysis, the numbers of death were summarized by ICD-10 chapters.

### Disease-to-disease distance matrices

Under the assumption that some diseases are more studied than others, we excluded diseases with the fewest features associated (first quartile) in each data dimensions to minimize the effects of low feature numbers on disease-to-disease comparisons. Summary statistics about the number of features in each dimension including the first quartile can be found in Table 1. After this trimming in each of the dimensions, we extracted the intersection of diseases among the six dimensions ending up with 514 diseases. Twelve diseases were further excluded since they had no disease name attached to the disease ID according to the annotation table retrieved from NCBI, thus reducing to a final set of 502 diseases.

**Table 1:** Summary statistics for number of features per disease.

|  | 1<sup>st</sup> Quartile | Median | Mean | 3<sup>rd</sup> Quartile |
| --- | --- | --- | --- | --- |
| # of associated genes | 12 | 51 | 313 | 192 |
| # of associated pathways | 6 | 12 | 57 | 42 |
| # of medicating chemicals | 2 | 4 | 14 | 12 |
| # of associated symptoms | 2 | 15 | 27 | 39 |
| # of chemicals targeting associated genes | 58 | 246 | 716 | 800 |
| # of associated chemicals | 5 | 36 | 198 | 125 |

#=number

For each data dimension, we calculated disease distances based on the associated features (genes, pathways, chemicals, and symptoms) using six different metrics in order to capture various aspects of similarity or dissimilarity in the data, providing a more robust and complete understanding of the relationships within datapoints compared to relying on a single metric: (1) Euclidean distance, (2) Hamming distance, (3) Cosine similarity, (4) Jaccard index, (5) Sørensen–Dice coefficient, and (6) overlap coefficient. For the latter three, vectors of features could be used as such, while for the first three metrics, binary vectors (associated yes/no) covering the union of all features were used. To compare and eventually merge the matrices generated by the six metrics, the Euclidean and Hamming distance matrices were normalized to a 0 to 1 standard scale by division by the maximum distance value. Cosine similarity, Jaccard index, Sørensen–Dice coefficient, and overlap coefficient, which originally measure similarities ranging from 1 to 0, were transformed into distance values ranging from 0 to 1 by subtracting the similarity value from 1. We aimed to combine the six distance matrices generated by the six metrics into one robust distance matrix reflecting the ultimate disease-to-disease distances in a particular data dimension. Therefore, we computed the Ipsen-Mikhailov distance among the six distance matrices using the netdist function of the R nettools package (version 1.1.0) and conducted agglomerative hierarchical clustering using the ward.D2 linkage criteria (*47*). Finally, a consensus distance matrix was computed by the stepwise averaging (arithmetic mean) of the six distance metrics according to branching points in the hierarchical clustering (Figure S3). By this, we generated one robust disease-to-disease distance matrix of 502 diseases for each of the six data dimensions. ICD-10-based disease similarity was calculated according to the formular suggested by Omura *et al.* (*48*).

### Comparison of dimensions and consensus score

To compute a consensus distance matrix for the 502 diseases based on the six data dimensions, we performed the same approach as when we combined the different distance metrics into one distance measure. Briefly, we computed the Ipsen-Mikhailov distance among the six data dimensions, conducted hierarchical clustering based on the Ipsen-Mikhailov distance and combined the six data dimensions by averaging according to their clustering in a hierarchical fashion (Figure 4, B). The resulting distance matrix contained the disease-to-disease consensus distance score that was used for subsequent analysis such as the clustering of diseases into groups.

### Clustering of diseases, visualization, and feature enrichment

Based on the disease-to-disease consensus distance score, agglomerative hierarchical clustering of the 502 diseases was performed using the ward.D2 linkage criteria. The optimal number of clusters was determined by the Dunn index as implemented in the dunn function of the clValid R package (version 0.7) (*49*). The Dunn index was computed for all numbers of clusters between 2 and 250. We observed a local maximum of the Dunn index for 71 to 78 clusters. The subsequent analysis was performed considering 71 clusters of diseases. We prompted ChatGPT3.5 (https://www.openai.com) to assign a suitable descriptive name for the 71 clusters. The prompt was: “Provide a meaningful and describing name or a describing category for the following list of diseases: [list of diseases]”. Subsequent analysis was focused on 16 candidate clusters containing at least five diseases. The remaining 55 clusters contained fewer than five diseases (n=21 clusters) and/or were an overall heterogenous mix of diseases, symptoms and generic phenotypes, as for example cellulitis, intestinal obstruction, cough, insomnia, and malnutrition. To visualize the distances among diseases in a two-dimensional space, we conducted multidimensional scaling on the disease-to-disease distance matrix using the cmdscale function in R. We plotted the coordinates of candidate disease clusters, while they had been calculated considering the complete distance matrix of 502 diseases. To identify overrepresented features (genes, pathways, symptoms, and chemicals) in each of the 71 clusters, we conducted a one-versus-all one-sided Fisher’s exact test per feature and cluster for each dimension with contingency matrices as shown in Table 2.

**Table 2:** Continency matrix for Fisher’s exact test to test feature overrepresentation.

|  | **Cluster j** | **All but cluster j** |
| --- | --- | --- |
| **Associated** | feature i associated with diseases in cluster j | feature i associated with diseases outside of cluster j |
| **Not associated** | feature i not associated with diseases in cluster j | feature i not associated with diseases outside of cluster j |

P-values were Bonferroni corrected for the number of clusters and the number of total features tested in each data dimension, *i.e.*, in case of the gene dimension, the adjusted p-value was calculated by nominal p-value*71(clusters)*21,020(genes). We further tested the differences between seven cancer clusters and nine non-cancer clusters by directly comparing cancer versus non-cancer clusters in a two-sided Fisher’s exact test. Pairwise Fisher’s exact test for three candidate clusters was performed comparing the frequency of features between two clusters directly.

We performed statistical analyses using the R statistical software. We performed visualization using R statistical software and biorender. To describe candidate gene lists on a functional level, we performed enrichment analysis with Enrichr (overrepresentation test) and visualisations with the KEGG Mapper (*50–52*).

## Acknowledgements

This work was supported by the EU Innovative Medicines Initiative 2 (IMI2) Biomap Project [grant agreement number 821511], Academy of Finland project UNICAST NANO [grant agreement number 322761], and the European Research Council (ERC) program, Consolidator project “ARCHIMEDES” [grant agreement number 101043848]. AS and AF were supported by Tampere Institute for Advanced Study (IAS).

## Conflict of Interest Disclosure

All authors declare no conflicts of interest.

## Data availability statement

All data were retrieved from public resources. Associated genes were retrieved from Open targets (https://www.opentargets.org/) (retrieved 15/02/2020). Associated pathways were retrieved from KEGG (https://www.genome.jp/kegg/) (retrieved 14/10/2021) and the Comparative Toxicogenomics Database (CTD) (https://ctdbase.org/) (retrieved 04/10/2019). Associated symptoms were retrieved from a study by Zhou *et al.* (*39*). Drug profiles were retrieved from Open targets (https://www.opentargets.org/) (retrieved 15/02/2020), LabeledIn (retrieved 11/02/2020), and a study from Wang *et al.* (*40*, *41*). Associated chemicals were retrieved from CTD (https://ctdbase.org/), SIDER (http://sideeffects.embl.de/), offsides and MEDI (retrieved 04/10/2019, 07/07/2020, 25/06/2020, and 02/07/2020, respectively) (*42*, *44*). Drug-target information was retrieved from DrukBank (https://go.drugbank.com/) (retrieved 11/06/2021), Open Targets (www.opentargets.org/) (retrieved 15/02/2020), and Pharos (https://pharos.nih.gov/) (retrieved 05/11/2020). Summary data for MedGen and PubMed identifiers (https://ftp.ncbi.nlm.nih.gov/pub/medgen/medgen_pubmed_lnk.txt.gz) (retrieved 13/04/2023), chemical identifiers and PubMed identifiers (https://ftp.ncbi.nlm.nih.gov/pubchem/Compound/Extras/CID-PMID.gz) (retrieved 18/04/2023), and Gene identifiers and PubMed identifiers (https://ftp.ncbi.nlm.nih.gov/gene/DATA/gene2pubmed.gz) (retrieved 14/02/2023) were retrieved from the ftp server of NCBI (https://ftp.ncbi.nlm.nih.gov/). Mortality data were retrieved from the Mortality database of the WHO (https://www.who.int/data/data-collection-tools/who-mortality-database, Mortality, ICD-10) (retrieved 03/10/23). The Supporting Data has been deposited in the Zenodo repository available at https://doi.org/10.5281/zenodo.10663498.

## Author contribution

**Lena Möbus:** Data curation, Formal analysis, Investigation, Validation, Methodology, Visualization, Writing - original draft, Writing - review & editing. **Angela Serra:** Formal analysis, Methodology, Supervision, Writing - review & editing. **Michele Fratello:** Data curation, Methodology, Writing - review & editing. **Alisa Pavel:** Data curation, Software, Writing - review & editing. **Antonio Federico:** Writing - review & editing. **Dario Greco:** Conceptualization, Supervision, Funding acquisition, Project administration, Resources, Writing - original draft, Writing - review & editing.

## Notes

https://doi.org/10.5281/zenodo.10663498

### Competing Interest Statement

The authors have declared no competing interest.

