## Supporting Information for "A Multi-Dimensional Approach to Map Disease Relationships Challenges Classical Disease Views"

### 1) Supporting Figures:

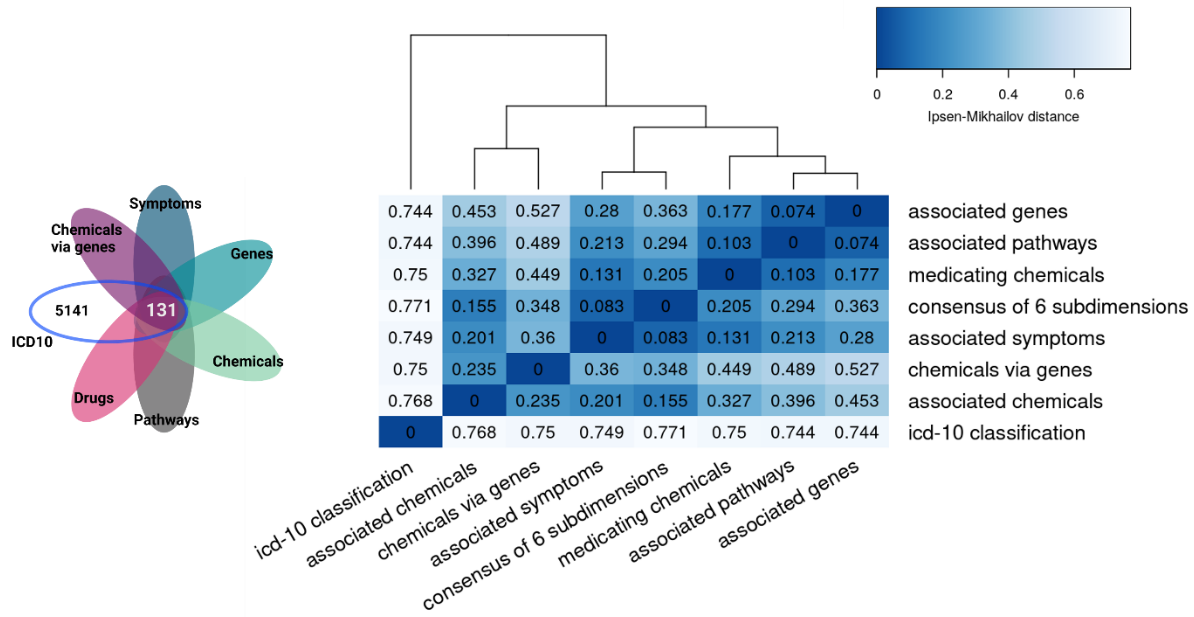


**Figure S1: Disease-to-disease distance matrices in six data dimensions and the ICD-10 system.** Left: The number of diseases with available ICD-10-MedGen match and the overlap with the six data dimensions. Right: Hierarchical clustering of the distance matrices in the six dimensions, the consensus distance matrix of them, and the ICD-10 distance matrix based on the Ipsen-Mikhailov distance computed among the matrices.

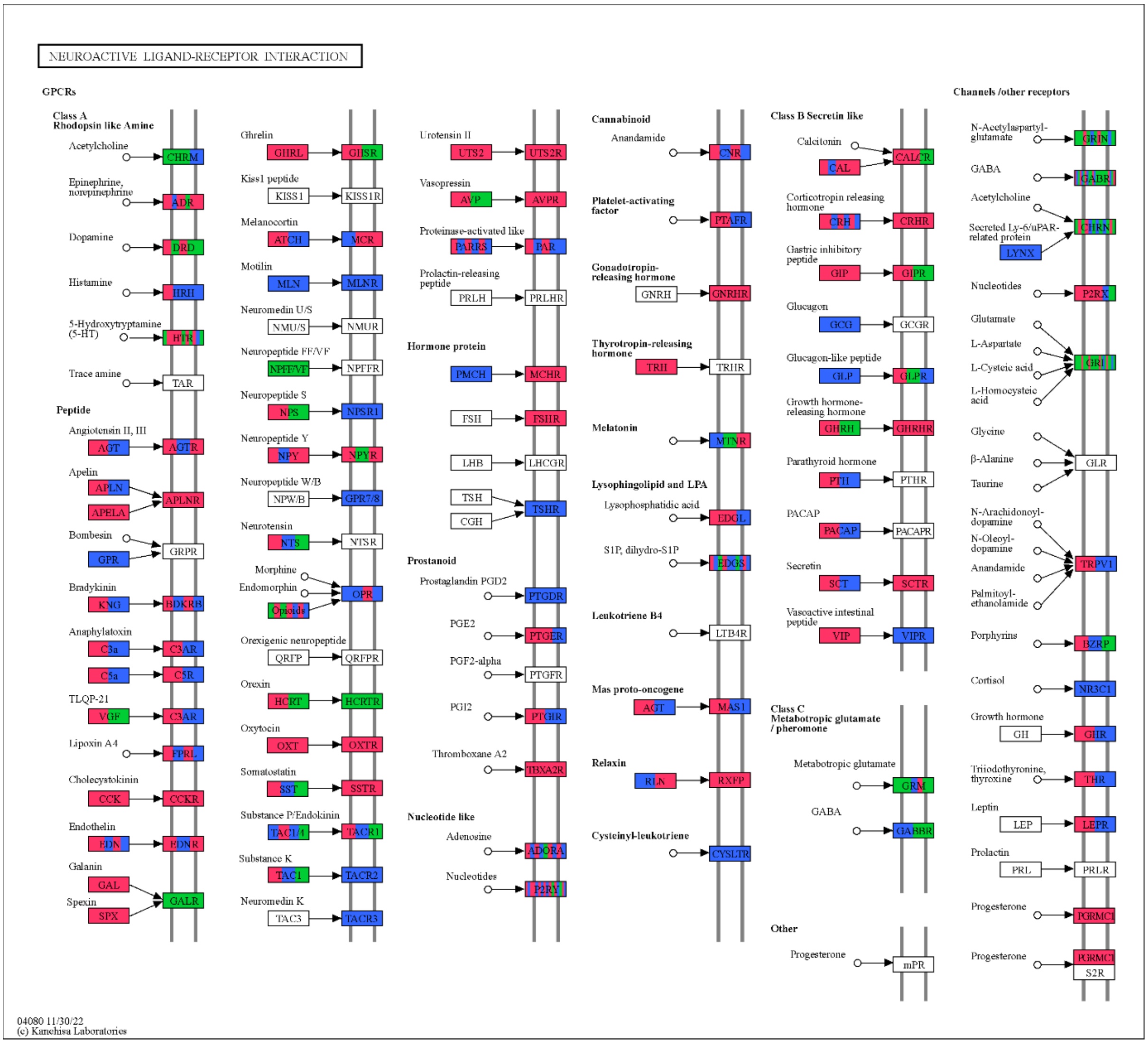


**Figure S2: Accumulation of enriched genes in neuroactive signaling.** The KEGG pathway of Neuroactive ligand-receptor interaction. Boxes represent units of gene products, and they are colored in green, red, and blue when their encoding genes were enriched in the cluster ”Neurological and Psychiatric Disorders”, "Cardiovascular and Metabolic Disorders”, and “Immunological and Respiratory Disorders”, respectively. This Figure was derived from the KEGG Mapper tool.

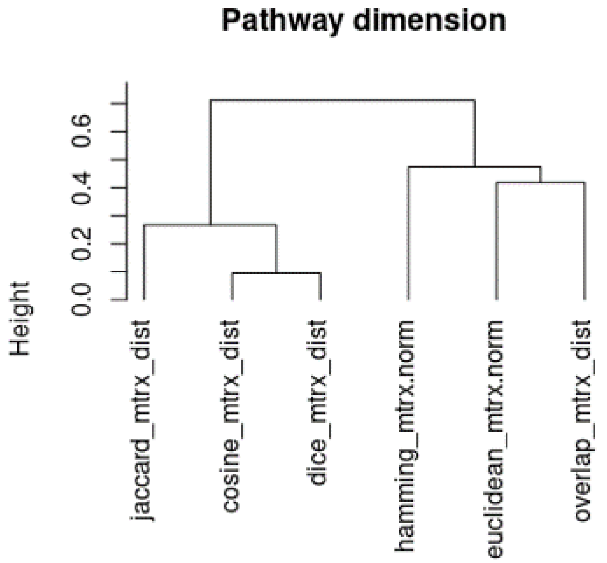


**Figure S3: Hierarchical clustering of six distance metrics for the pathway dimension.** To combine the six distance metrics into one robust distance measure, we took the mean of the cosine and dice distance matrix. The resulting value was averaged with the Jaccard distance matrix resulting in the branch-1 matrix. The mean of the Euclidean matrix and the Overlap matrix was taken. The resulting matrix was averaged with the Hamming matrix resulting in the branch-2 matrix. The mean of the branch-1 matrix and the branch-2 matrix yielded the final distance matrix for the pathway dimension.

### 2) Supporting Tables

**(available at Zenodo https://doi.org/10.5281/zenodo.10663498)**

Table S1: Number of associated diseases and number of linked PMIDs for each feature in the six data dimensions

Table S2: Cluster assignment and further information for 502 diseases

### 3) Supporting Data

**(available at Zenodo https://doi.org/10.5281/zenodo.10663498)**

Supporting Data 1: Data_diseases_of_six_dimensions.RData

Summary data for the six data dimensions.

Supporting Data 2: Consensus_distance_matrices_six_dimensions_and_icd10.RData

The consensus distance matrices for six dimensions and the distance matrix of ICD-10 codes matching with a MedGen IDs.

Supporting Data 3: Enriched_features_one_vs_all_clusters_six_dimensions.RData

Enriched features of six data dimensions by Fisher's exact test for 71 clusters of diseases.

Supporting Data 4: Enriched_features_cancer_vs_noncancer_clusters_six_dimensions.RData

Enriched features of six data dimensions by Fisher's exact test for seven cancer clusters versus nine non-cancer clusters of diseases.

Supporting Data 5: Enriched_features_pairwise_comparisons_of_3_candidate_clusters_six_dimensions.RData

Enriched features of six data dimensions by Fisher's exact test for three candidate clusters of diseases. ("Neurological and Psychiatric Disorders", "Cardiovascular and Metabolic Disorders", and "Immunological and Respiratory Disorders").
